# Neuro-genetic plasticity of *Caenorhabditis elegans* behavioral thermal tolerance

**DOI:** 10.1101/583120

**Authors:** Gregory W. Stegeman, Denise Medina, Asher D. Cutter, William S. Ryu

**Affiliations:** University of Toronto Department of Ecology and Evolutionary Biology, Toronto, Ontario M5S3B2 Canada; University of Toronto Department of Physics, Toronto, Ontario M5S3B2 Canada; Donnelly Centre University of Toronto, Toronto, Ontario M5S3B2 Canada

**Keywords:** *C. elegans* behavior, thermal ecology, computational ethology

## Abstract

**Background:** Animal responses to thermal stimuli involve intricate contributions of genetics, neurobiology and physiology, with temperature variation providing a pervasive environmental factor for natural selection. Thermal behavior thus exemplifies a dynamic trait that requires non-trivial phenotypic summaries to appropriately capture the trait in response to a changing environment. To characterize the deterministic and plastic components of thermal responses, we developed a novel micro-droplet assay of nematode behavior that permits information-dense summaries of dynamic behavioral phenotypes as reaction norms in response to increasing temperature (thermal tolerance curves, TTC).

**Results:** We found that *C. elegans* TTCs shift predictably with rearing conditions and developmental stage, with significant differences between distinct wildtype genetic backgrounds. Moreover, after screening TTCs for 58 *C. elegans* genetic mutant strains, we determined that genes affecting thermosensation, including *cmk-1* and *tax-4*, potentially play important roles in the behavioral control of locomotion at high temperature, implicating neural decision-making in TTC shape rather than just generalized physiological limits. However, expression of the transient receptor potential ion channel TRPA-1 in the nervous system is not sufficient to rescue rearing-dependent plasticity in TTCs conferred by normal expression of this gene, indicating instead a role for intestinal signaling involving TRPA-1 in the adaptive plasticity of thermal performance.

**Conclusions:** These results implicate nervous system and non-nervous system contributions to behavior, in addition to basic cellular physiology, as key mediators of evolutionary responses to selection from temperature variation in nature.

## Background

Behaviors are the primary way that animals interact with their environments and, in so doing, connect genetics and physiology with ecology. Natural selection favours those alleles of genes that allow animals to sense stimuli and react in ways that maximize fitness (1, 2). While many aspects of behavior are heritable, it remains an ongoing challenge to discover mechanisms of when and where genes will influence specific aspects of behavior (3). Behaviors as phenotypes are inherently complex because of the combination of environmental and genetic factors that influence them, ranging from internal and external conditions of an individual, to widespread functional pleiotropy and gene by environment interactions, to previous experience of individuals and their social interactions (4, 5). Because many behavioral responses are reactions to stimuli, it is natural to describe a behavioral trait as a functional response, the norm of reaction to a range of environmental inputs (6, 7). Temperature provides a ubiquitous environmental input that is particularly crucial to organismal fitness and to behavior, commonly characterized phenotypically as thermal performance curves (8-10). Behavior is the primary strategy of ectothermic animals to regulate their body temperatures, with sensing, orienting and navigating the thermal landscape being critical for fitness (11, 12). Invention of quantitative behavioral assays has helped geneticists unravel the roles of genes in behavior by controlling as many variables as possible and by quantifying behavioral traits as accurately as possible (13). In order to quantify the dynamic temperature-dependent behavior of *Caenorhabditis* nematode and to decipher its neuro-genetic control, we developed a novel assay to quantify locomotory behavior through a range of ecologically relevant temperature.

As small-bodied ectotherms, *C. elegans* worms regulate their body temperature through movement, making the sensing and orienting to thermal stimuli crucial for survival and reproduction in nature. *C. elegans* navigates it thermal landscape predominately through two behaviors: thermotaxis (14) and thermal avoidance (15, 16). Although much is known about the molecular and cellular mechanisms of thermosensation and thermosensory neural circuits in *C. elegans* (17), much less is known about the general adaptive mechanisms that *C. elegans* uses as it explores its entire ecological temperature range. Powerful approaches to study temperature-dependent behaviors in *C. elegans* include “classic” assays in Petri dishes as well as automated and microfluidic measurement (18-20), but none of these assays have characterized *C. elegans* thermal performance curves. Our goals led us to develop a new assay for *C. elegans* temperature-dependent behavior to quantify thermal tolerance from video recordings, where we capture the behavioral responses by a given genotype across a range of temperatures as a ‘norm of reaction’.

By framing the problem of understanding behavior as a set of thermal reaction norms in the genetically-tractable *C. elegans* system, we aimed to answer key questions about deterministic and plastic contributors to behavior. How do genes and sensory inputs modulate behavioral thermal performance? Do neural decisions or fundamental physiological limitations drive behavioral performance at high temperatures? How sensitive is late-life thermal performance to early-life experience and can genes modulate that sensitivity? In addressing these outstanding questions, we aim to decipher the links between thermal ecology and the neural and genetic controls over complex behavioral responses to stimuli.

After constructing and implementing a micro-droplet assay device for locomotory thermal performance in *C. elegans*, we demonstrate its efficacy for quantifying behavioral differences among natural isolates and a panel of gene mutants. By quantifying the reaction norm of the behavioral response to precisely controlled changes in temperature (thermal tolerance curve, TTC), these experiments provide evidence for a neural “decision” by animals to stop locomotion as temperature increases. Animals with genetic defects in thermosensation shift the decision-making process to continue swimming at higher than normal temperatures or to cease moving at lower than normal temperatures. Therefore, the declining locomotory behavior with increases in temperature does not solely reflect high-temperature physiological limits, as usually presumed for other organisms (10). Plasticity in animal TTC profiles derives from differences in age and rearing conditions. We also determined that non-neural regulation of TTCs is important, particularly for plasticity of TTC profiles in response to larval rearing conditions, with intestinal expression of the *trpa-1* ion channel implicated in this process.

## Results

### *C. elegans* swimming behavior as a thermal reaction norm

We developed a micro-droplet assay of nematode worm swimming behavior that permits relatively high-throughput and information-dense quantification of individual locomotion in response to precise temperature manipulation. We then characterized animal movement as norms of reaction, finding that nematode behavior follows the pattern of a classic thermal performance curve in the high temperature range (12, 21) with locomotion most rapid at benign temperatures and steeply dropping off in performance as temperature increases until reaching paralysis (Fig. 1A). By contrast, constant benign temperatures yield much smaller, albeit significant, changes of motility over time through the course of a 21 min. assay (Fig. S2B) (repeated measures ANOVA F_19,7_=4.21, p<0.0001). Worms are slightly more active when they experience a constant 25°C compared to 21°C or 23°C (repeated measures ANOVA F_2,92_=7.96, P=0.0006; also apparent in the dynamic TPC profile, Fig. S2B).

**Figure 1.**
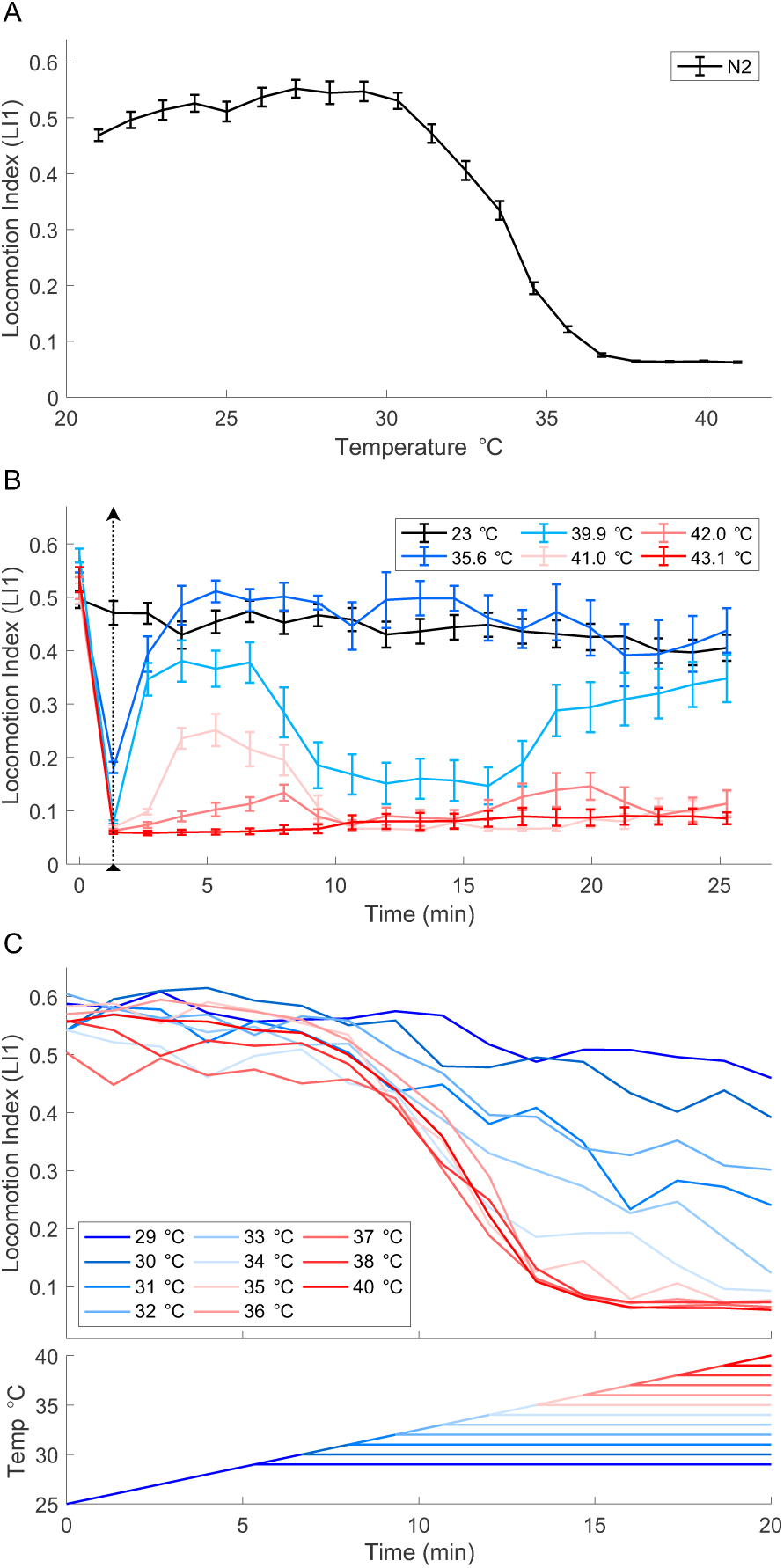
Thermal tolerance curve for wildtype *C. elegans*. (A) *C. elegans* thermal tolerance curve. Locomotion index (LI1) with temperature increments from 21°C to 41°C. Wildtype N2 adult hermaphrodite animals were reared at 23°C. 54 worms and ≥35 animals retained in calculations at each temperature step. (B) *C. elegans* locomotion index profiles in response to acute high temperature exposure from baseline 23°C (black arrow dotted line, 80 sec exposure). Error bars indicate ± SEM. 54 individuals tested per series; ≥34 retained in calculations at each step. (C) Locomotion index over time at constant hold temperature after incrementing up from an initial 25°C (upper half of panel). Hold temperature shown on the bottom half of the panel, with hold temperature of 40°C comparable to the standard TPC assay; 18 worms tested per series, ≥9 worms included in LI calculations at each step.

Temperature-induced paralysis is reversible and the dynamics of recovery happen at multiple time scales. We tested reversibility by submitting worms to brief 80 sec, high temperature exposures (35°C-43°C). During exposure to temperatures >39°C, all worms quickly ceased movement (Fig. 1B), but when returned to a benign temperature (23°C, rearing temperature), worms recovered in a manner inversely related to the exposure temperature. Worms exposed to 35.6°C rapidly recovered to control activity but those exposed to 43.1°C did not recover at all. For intervening exposure temperatures, worms surprisingly showed non-monotonic recovery with an early phase partial or weak recovery and then a second phase of recovery later in the assay (Fig. 1D). However when we extended the room temperature observation to 77 minutes post-exposure, we observed that animals exposed to 41.0°C and 42.0°C which recovered within a few minutes, ultimately ceased locomotion after about an hour (Fig. S2).

Thermal paralysis can be induced over time at temperatures that are below the thermal paralysis threshold. To test sub-threshold temperatures, we increased temperature incrementally from 25°C as before, but then held it constant for the remainder of the assay at various elevated temperatures. We found that worm locomotion declines toward immobility for any ‘hold temperature’ above 35°C (Fig. 1C). Even though worms swim at the start of the 35.6°C step, their eventual paralysis does not require that they experience higher temperatures. In fact, worms exposed to hold temperatures of ∼33°C also come close to paralysis within the 21 min. assay, whereas hold temperatures less than ∼30°C exert at most a minimal effect on locomotion (Fig. 1C). These experiments implicate a cumulative effect of temperature exposure above a critical threshold of ∼30°C, and a delay in the ultimate outcome of exposure to temperatures higher than ∼33°C. In agreement with our work, previous research on heat shock showed that a 20 min. exposure to 40°C led to *C. elegans* quiescence, a sleep-like state, lasting several hours (22) and < 25% of worms survived a 15 min. exposure to 39°C (23). These results show that the TTC is a complicated convolution of thermal thresholds and cumulative effects, and so the precise shape of the TTC depends on the time spent at each target temperature.

### Plasticity of *C. elegans* behavioral thermal tolerance curves: rearing conditions and life stage

In order to place the micro-droplet thermal performance assay in a broader ecological context, it is important to consider other life stages and rearing conditions. Given how strongly rearing temperature affects growth rates and other aspects of *C. elegans* biology, as well as the role of thermal acclimation and memory in thermotaxis and isothermal tracking behaviors (24-27), we expected rearing temperature to affect thermal tolerance curves of locomotory behavior. We quantified this sensitivity by comparing N2 worms reared at 15°C, 20°C, 23°C and 25°C in terms of their TTCs as the animals experienced temperature increments from 21°C to 40°C (Fig. 2A). We found that cooler rearing temperatures led to more rapid declines in swimming behavior as we raised the temperature (Fig. 2A). Opposing this trend, however, *C. elegans* reared at 25°C exhibit reduced locomotion compared to worms reared at 23°C. We hypothesize that animals reared at 25°C may have a developmental fate difference that alters the TTC because this higher temperature is known to compromise *C. elegans* fecundity (28, 29).

**Figure 2.**
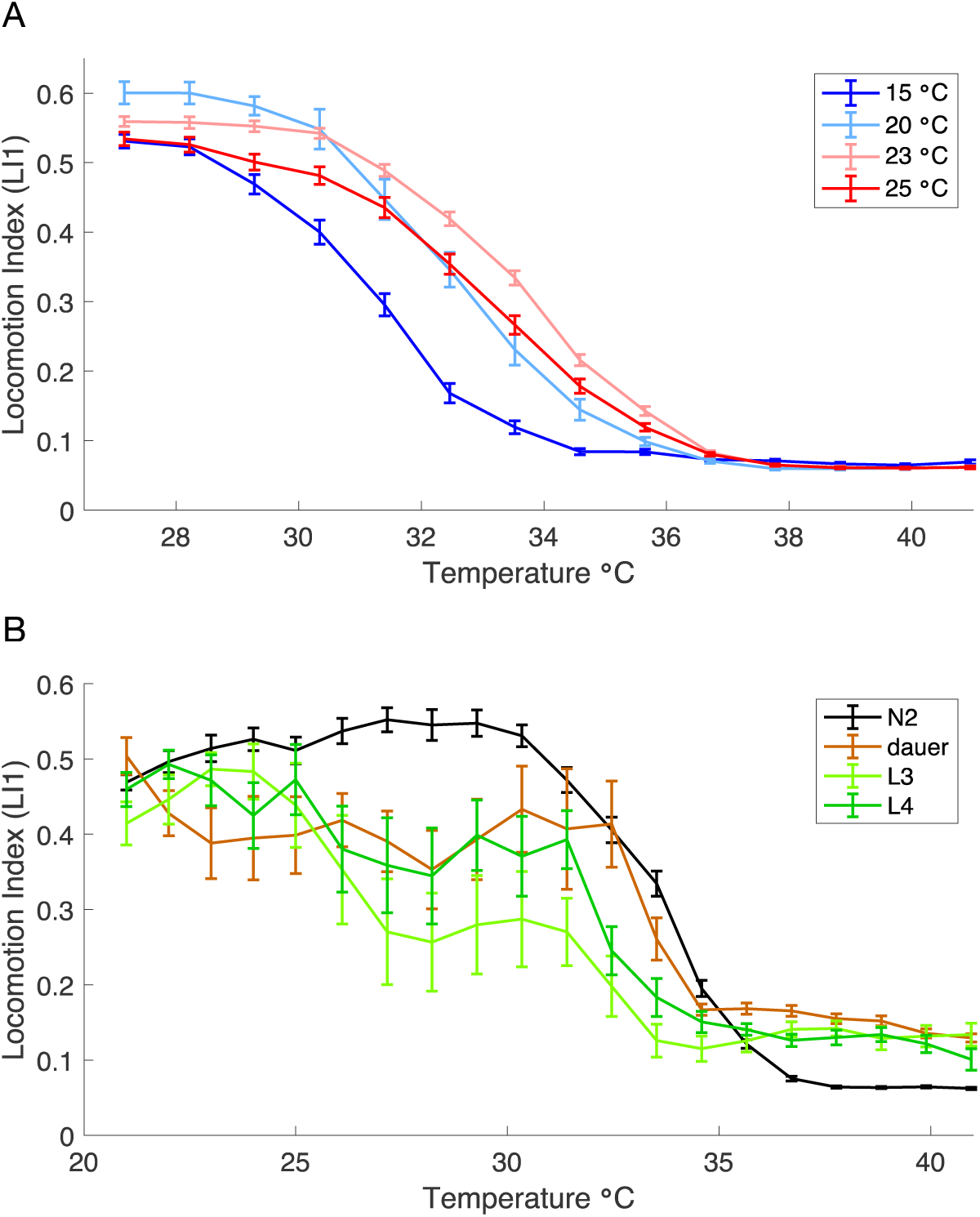
Effect of rearing temperature and developmental stage on *C. elegans* TPCs. (A) Rearing conditions at both high and low thermal extremes shift wildtype N2 *C. elegans* thermal performance curves relative to rearing at intermediate thermal conditions. 36-114 animals tested per thermal rearing condition, ≥16 individuals included in calculations at each assay step. (B) TPCs for adult and larval developmental stages. 9-27 worms tested per treatment, ≥5 worms included in calculations at each assay step. Error bars indicate ± SEM.

We also contrasted adult TTCs with those of larvae and found that animals in the two final larval stages (L3 and L4) slow their locomotion more readily than do adults in response to increasing temperatures, even at relatively benign temperature points (∼26°C; F_2,62_=8.49, P=0.0006, Dunnett’s post-hoc test L3 P=0.0021, L4 P=0.0072; Figure 2B). The L3 animals tend to move on average more slowly than L4 animals and essentially stop movement above 33°C, whereas L4 worms continue some movement up to 35.6°C (Figure 2B). By contrast, dauer larvae tend to move at a slow and consistent pace at benign temperatures up to ∼32°C when they start to slow at a rate similar to adult animals (Fig. 2B). Dauer larvae are a long-lived stress resistant alternate larval pathway for many nematode species including *C. elegans*, capable of surviving higher temperatures than normal larvae (30, 31). These experiments point to different developmental stages as having distinct TTC shapes, perhaps reflecting different optimal strategies for actively developing animals, distinct neural processing circuitry, or different capability of handling temperature stress.

### Genetic perturbation of *C. elegans* behavioral thermal tolerance curves

Many *C. elegans* genes affect behaviors, so we screened 58 gene mutant strains to see how these genes might change behavioral thermal performance compared to two “wildtype” control strains in the micro-droplet assay, with temperature incrementing from 27°C to 40°C. In particular, we aimed to analyze mutants with known phenotypic disruptions of thermotaxis, thermosensation, mechanosensation, chemosensation, locomotion, and neural function (Fig. 3 and S3). The Hawaiian wildtype strain (CB4856) appears to slow movement more sharply above 33°C than the N2 (Bristol) strain, but nevertheless continues locomotion to the same high temperatures as N2, with comparable locomotion index scores above 36°C (Fig. 3B). The TTC for the *npr-1* mutant performs similar to CB4856 (Fig. 3B), which has a functionally similar allele of *npr-1* (32). Most mutant strains exhibited lower locomotion indices at high temperatures compared to the wildtype N2 (F_59,1550_=10.79, P<0.0001, 47 mutant strains lower and only *cmk-1* higher than wildtype with Dunnett’s post-hoc tests P<0.05 at 35.6°C; Fig. 3), including the other wildtype strain (N2 LI1=0.56 vs. CB4856 LI1=0.48 at 27.2°C). Indeed, about half of the mutants show very low activity even under benign conditions, consistent with them having basic roles in locomotion. In general, it is difficult to dissect the mechanistic implications of strains with very poor locomotion because we cannot easily distinguish general locomotory defects from those caused by perturbation specific to thermal paralysis. However, we can use the condition of thermal paralysis to compare curves, even for those with initial low activity. While most curves share the same shape as the wild-type TTC, some curves reach paralysis at a much higher temperature (*dgk-1, goa-1, mec-6, tax-4, unc-46*; F_59,1540_=9.64, P<0.0001, Dunnett’s post-hoc tests P<0.02 at 37.8°C) (Fig. 3B) or lower temperature (*mec-3, odr-1, odr-3, pkc-1, daf-21, hsf-1,* all Dunnett’s post-hoc test P<0.0001 at 35.6°C) (Fig. 3C), suggesting non-trivial perturbations affecting the TTC. And finally, we had hypothesized that a number of mutants would have strongly altered TTCs, but their TTCs were relatively unchanged compared to the WT (e.g. *hsf-16.48*) (Fig. 3B).

**Figure 3.**
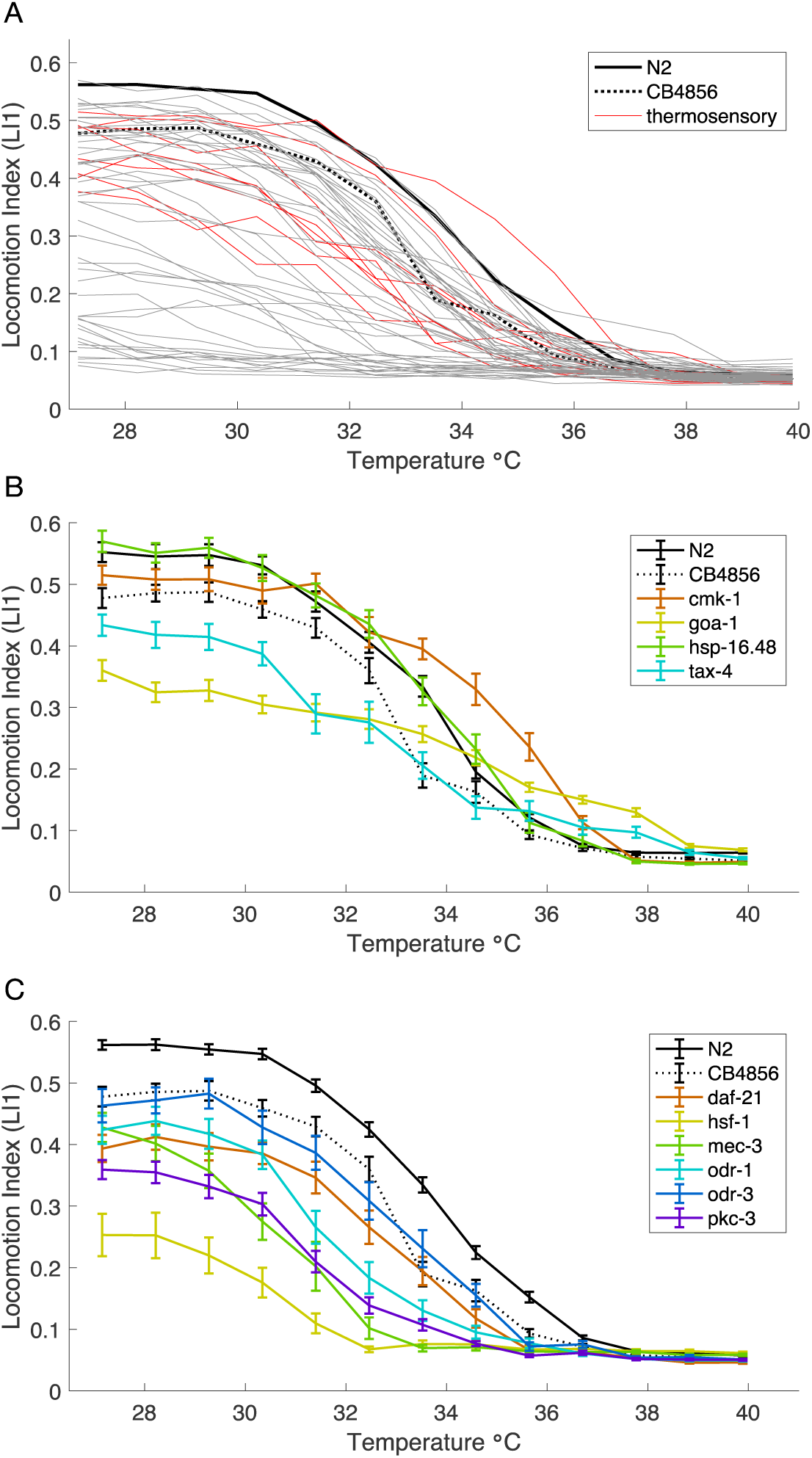
Effect of mutations on *C. elegans* TPCs. (A) Thermal reaction norms of swimming behavior for 58 *C. elegans* genetic mutant strains (gray and red lines) and two wildtype strains (black line = N2, dashed line = CB4856). All worms reared at 23°C; 17 - 36 individuals included in calculations at each assay step for each strain (Supplementary Table S1). TPCs for subsets of mutant strains with sensory disruptions from (A) are shown for strains with similar or higher paralysis threshold temperatures than wild-type (B) and strains with lower paralysis temperatures than wild-type (C).

### Neuronal gene disruptions can enhance or reduce TTC responses

The eight thermosensory mutants provide a class of mutations of special interest (Fig. 3, Fig. S3B). Might a neural “decision” act to stop locomotion as temperature increases? We hypothesized that if neural control drives the slowing of locomotion at higher temperatures, then thermosensory mutant strains should exhibit distinct TTCs compared to wildtype and implicate sensation or perception of temperature in the response rather than some fundamental physiological limit.

Indeed, we found that two thermosensory mutant genotypes outperform the wildtype over part of the thermal tolerance curve [*cmk-1*(oy21) at 34.6°C F_59,1563_=14.39, P<0.0001, Dunnett’s post-hoc test P<0.0001; *tax-4*(p678) at 37.8°C as above F_59,1540_=9.64, P<0.0001, Dunnett’s post-hoc tests P<0.0001; Fig. 3B]. However, *cmk-1*(oy21) and *tax-4*(p678) differ in the trend of how they manifest higher locomotion than wildtype at high temperatures. The *cmk-1*(oy21) TTC is shifted toward hot temperatures whereas the *tax-4*(p678) TTC shows a nearly linear decay from benign temperatures that eventually yields higher locomotion than N2 from 36°C-39°C (Fig. 3B). Interestingly, an alternative allele of *tax-4*(ks11) that is a missense rather than a nonsense mutation (33) does not show elevated movement at high temperatures like the p678 allele (Dunnett’s post-hoc test P=0.12 for *tax-4*(ks11) at 37.8°C) (Fig. S3B). Both of these loci disrupt development of AFD, the main thermosensory neuron in *C. elegans* (34). The fact that these AFD-expressed thermosensory genes (*cmk-1, tax-4*) both disrupt TTCs, either positively or negatively, implicates neuronal thermal sensory measurement as important in the thermal locomotory performance and indicates that reduced high-temperature locomotion is not simply a result of general physiological limits.

Given the neural connection to TTCs, we explored whether mutations that alter neural circuitry or sensory structures that impact mechanosensation and chemosensation might also disrupt TTCs. Of these 22 mutants, five became paralyzed at lower temperatures than N2 (*mec-3, odr-1, odr-3, pkc-1, daf-21*; all Dunnett’s post-hoc test P<0.0001 at 35.6°C) (Fig. 3C). The other 18 strains exhibited deficits in thermal performance at moderately high temperatures (32°C −36°C), but most were comparable in TTC profile to the CB4856 Hawaiian wildtype strain (7/11 mechanosensory and 10/11 chemosensory mutants lower LI1 than N2 at 35.6°C; Fig. S3C-D). No mutants from these sensory classes significantly enhanced locomotory thermal performance at high temperatures, though two trended that way (*age-1* at 36.7°C, involved in chemosensation; *glr-1* at 37.8°C, involved in mechanosensation), and 7 of the remaining 20 mutants showed severely compromised TTCs (Fig. S3C-D). These mutants show that neuro-modulation of thermal performance is not limited to changes in thermosensory genes and cells. Given the limited number of neurons in *C. elegans* (just 302 neurons in adult hermaphrodites, (35)), many circuits are used for multiple purposes. Even individual sensory neurons may sense temperature as well as other stimuli (36). The influence of chemo-and mechanosensory genes on TTCs suggests that pleiotropic effects of neuronal disruption might work in the same or parallel circuits to those affected by thermosensory mutants.

Our mutant screen also included six genes involved in dopamine signaling, which is necessary for arousal and activity in *C. elegans* (37). In other studies, *dop-1* mutants usually have reduced locomotory levels (38), which we also observed (Dunnett’s post-hoc test P<0.0001 at 35.6°C; Fig. S3E). Other dopamine pathway mutants showed TTCs similar to the CB4856 Hawaiian wild isolate, which shows reduced locomotory activity at moderately high temperatures (33°C-36°C) compared to N2 (3/6 dopamine mutants lower LI1 than wt at 35.6°C with Dunnett post-hoc test P<0.004; Fig. S3E). Therefore, the changes in locomotory activity with increasing temperature may not involve a major role of dopamine signaling. However, the *goa-1* mutant is known for being hyperactive (39), and in our experiment it maintains much faster locomotion than wildtype in the region of the TTC above 35°C and has relatively high locomotion even at 37.8°C (Dunnett’s post-hoc test P<0.0001 at 37.8°C; Fig. 3B), where N2 is nearly motionless. GOA-1 is part of heterotrimeric G-protein alpha subunit G_o_ and, as such, interacts genetically with many other pathways thus making it very pleiotropic, especially with behavior phenotypes. The enhanced activity of *goa-1* mutant animals at high temperatures reinforces the idea that the wildtype TTC is not solely a by-product of physiological limits imposed by high temperatures (e.g. muscle function) but reflects in part behavioral decision-making by the animal to modulate locomotory activity.

To explore the role of the heat shock response, we tested two heat shock protein mutants (Fig 3, Fig. S3F). Only one of two mutants affecting the heat shock response in our screen exhibited an atypical TTC: the *hsf-1* mutant showed a locomotion deficit even at relatively benign temperatures (27.2°C Dunnett’s post-hoc test P<0.0001) but became immobile by ∼32°C, an exceptionally low paralysis temperature (Fig. 3C, 32.5°C Dunnett’s post-hoc test P<0.0001). Note that HSP20 chaperone *hsp-16.48* phenotype was indistinguishable from wildtype (Fig. 3B, Dunnett’s post-hoc test P=1.0 at 27.2°C and P=0.55 at 37.8°C). Even if triggered, however, expression of heat shock pathways may not fully manifest in the timeframe of the assay (40, 41), though pre-conditioning animals to heat stress in advance of assessing TTCs could test for more active roles of heat shock response in behavior as for survival (23).

### Non-neural *trpa-1*-mediated behavioral plasticity in response to rearing temperature

The TRPA-1 transient receptor potential ion channel activates at lower environmental temperatures, signaling for changes in gene expression that modulate lifespan in *C. elegans* (42). Might *trpa-1* also be important in regulating behavioral thermal performance in some way? It is a cold sensitive channel and is involved in thermosensation and mechanosensation in *C. elegans* (43), but it is also broadly expressed in non-neuronal tissues including the intestine. The TRPA1 family of channels also detect reactive electrophiles and acidification and other nociceptive stimuli (44). Under our standard locomotion assay conditions (benign 20°C rearing temperature), the *trpa-1*(ok999) knockout mutant strain exhibits modestly reduced locomotory activity compared to the N2 wildtype with equivalent baseline motility. To assess whether this cold-sensing channel might play a deeper role in behavioral thermal performance, we manipulated rearing temperatures of mutant and wildtype *trpa-1* alleles and contrasted their TTCs.

We hypothesized that if the downstream targets for *trpa-1* signaling affect acclimation behavior, as well as longevity, then *trpa-1* knockout animals might show a distinctive sensitivity to rearing conditions. Comparison of TTCs for *trpa-1*(ok999) in response to different rearing temperatures (15°C, 20°C, 25°C) revealed that it indeed shows reduced TTC plasticity across rearing conditions (rearing temperature differences for wt at 32.5°C F_2,147_=36.8, P<0.0001; for *trpa-1*(ok999) at ∼32C at 32.5°C F_2,145_=0.91, P=0.41; Fig. 4A-B; Fig. S3, S4). While TTCs for wildtype animals shift toward colder temperatures when reared cold and shift warmer when reared warm, the TTC for the *trpa-1* mutant strain remains nearly unchanged regardless of rearing temperature (Fig. 4A-B). In other words, knockout of *trpa-1* canalizes the TTC, making its shape largely insensitive to rearing conditions. A consequence of the *trpa-1* insensitivity to rearing conditions is that *trpa-1*(ok999) animals exhibit enhanced locomotion compared to N2 at warmer assay temperatures when they are both reared at 15°C; only rarely do mutants “outperform” the wildtype genotype in our screen of genetic perturbations.

**Figure 4.**
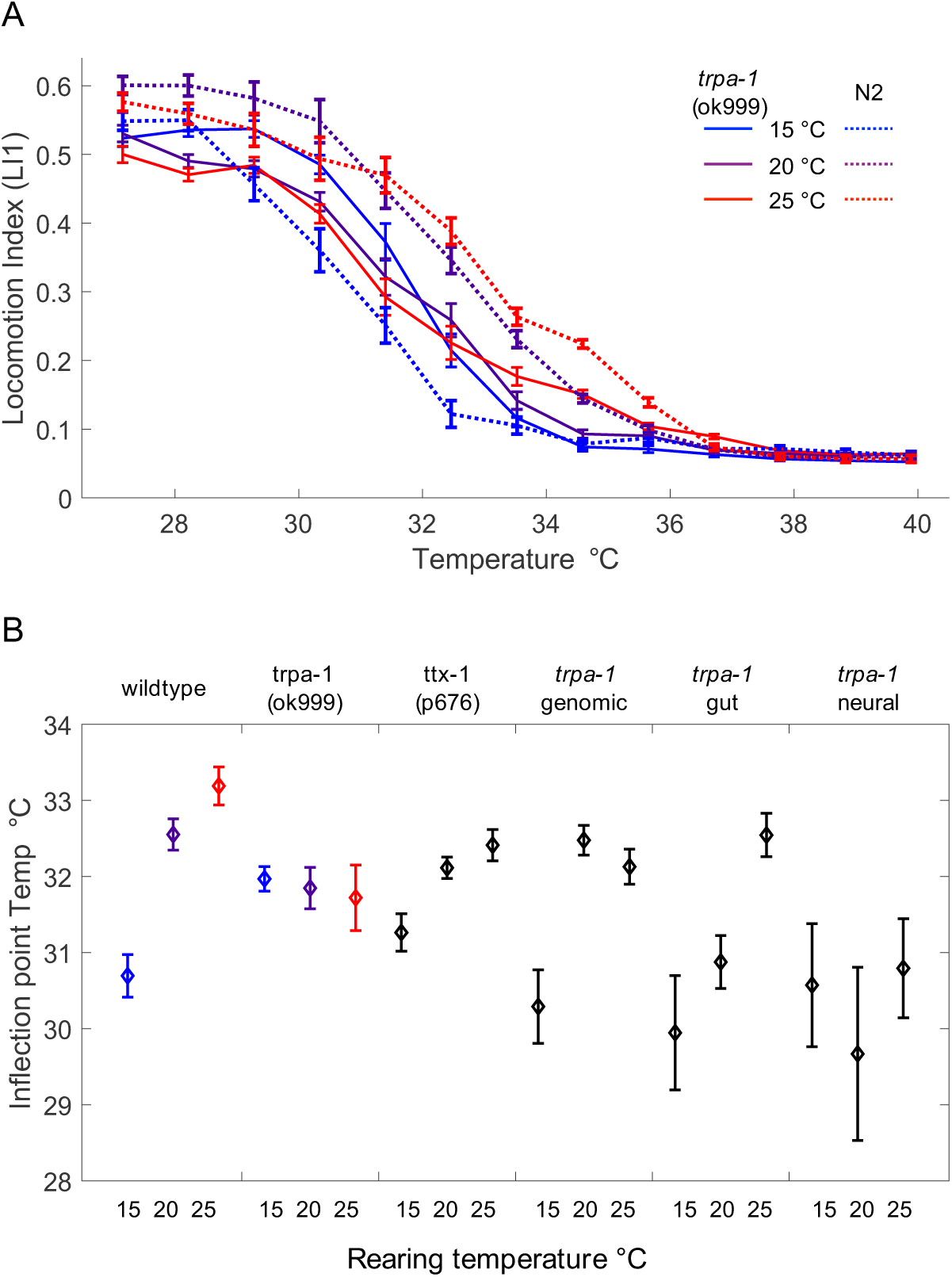
Comparison of wildtype and *trpa-1* mutants over different rearing temperatures. (A) The sensitivity of the locomotion index TTC to rearing temperature observed for wildtype N2 *C. elegans* (pale lines) is reduced for *trpa-1* mutant animals (dark lines), which show similar TTCs regardless of rearing conditions. Error bars indicate ±1SEM; 54 animals tested per treatment, ≥44 individuals included in calculations at each assay step. (B) Comparison of inflection point estimates for locomotion index TTC curves for wildtype N2, *trpa-1* and *ttx-1* mutants, and *trpa-1* expression-rescue strains (see Table 1 and Fig. S3). Similar inflection point values for different rearing conditions of a given genotype indicate low TTC plasticity; distinct values for different rearing conditions indicate high TTC plasticity. Genomic and gut expression-rescue constructs for *trpa-1* also rescue TTC plasticity, unlike the neural expression-rescue construct. Values for wildtype N2 and *trpa-1(ok999)* correspond to TTCs shown in (A). Error bars indicate 95% confidence intervals from a three-parameter logistic function fit to locomotion index (LI1) values for each strain and rearing temperature; 45-54 animals tested per treatment, ≥31 individuals included in calculations at each assay step.

**Table 1.** *trpa-1* construct lines assayed for TTCs.

| Strain* | Construct | Genetics | Explanation |
| --- | --- | --- | --- |
| RB1052 | - | *trpa-1(ok999)* | *trpa-1* knockout mutant |
| TQ1643 | XuEx601 | *XuEx601[Pges-1::trpa-1::SL2::yfp + Punc-122::DsRed]* | gut promotor overexpression |
| TQ1648 | XuEx606 | *XuEx606[Prgef-1::trpa-1::SL2::yfp + Punc-122::DsRed]* | neural promotor over expression |
| TQ2706 | XuEx619<br>&<br>XuEx735 | *trpa-1(ok999); lin-15; xuEx735[Pges-1::trpa-1::SL2::mcherry2 + Punc-122delta::yfp]; xuEx19[Plfe-2L::GCaMP1.3 + Plfe-2L::DsRed2b + lin-15(+)]* | *trpa-1* rescued with own genomic DNA promotor |

\*Xiao et al. (2013); We tested strain TQ2706 as a *trpa-1* rescue using our own upstream genomic promotor. Lines with constructs XuEx601 and XuEx606 were backcrossed separately to strain RB1052 to create tissue-specific *trpa-1* rescue lines, confirmed by PCR and fluorescence.

These results are consistent with the hypothesis that *trpa-1* mutants do not adjust their behavior normally to rearing temperature. Previous work found that overexpression of TRPA-1 in neurons and the gut led to longer lifespans at 20°C, with the gut expression exerting a greater effect (42). Because we demonstrated that TTCs are at least partly affected by thermosensory neural circuitry, we hypothesized that TRPA-1 expression in neurons might also be important for the mutant phenotype. We therefore next quantified the TTCs for *trpa-1* expression rescue lines in a *trpa-1* null genetic background, using transgenic constructs with its endogenous promoter or using tissue-specific rescue of expression with neural or gut *trpa-1* overexpression lines.

Expressing *trpa-1* with its genomic promotor yields a TTC with more plasticity than the gene knockout in response to rearing temperature, especially for cool 15°C rearing, indicating at least partial rescue of rearing-dependent TTC plasticity (rearing temperature differences for *trpa-1* genomic rescue at 32.5°C F_2,140_=17.37, P<0.0001; Fig. 4B; Fig. S3, S4). Gut-specific expression of *trpa-1* also exhibited nearly normal plasticity from rearing temperature, especially for warm 25°C rearing (rearing temperature differences for *trpa-1* gut rescue at 32.5°C F_2,127_=18.41, P<0.0001; Fig. 4B; Fig. S3, S4). The imperfect restoration of TTC plasticity might result from over-expression of *trpa-1* or the fluorescent markers in the transgenic constructs of the rescue lines that could adversely affect animal health. Interestingly, and counter to our initial predictions, neural-specific *trpa-1* expression did not restore any plasticity to the TTCs in response to rearing conditions (rearing temperature differences for *trpa-1* neural rescue at 32.5°C F_2,136_=0.90, P=0.41; Fig. 4B; Fig. S3, S4), and the TTCs were shifted toward more rapid decline in locomotory activity with ambient temperature changes than for any of the other *trpa-1* experimental lines (Fig. 4B; Fig. S3). Together, these experiments imply that the downstream effects of TRPA-1 thermosensory activity derive from intestinal expression during development, and perhaps expression in other tissues conferred by the endogenous promoter, but not from TRPA-1 thermosensory signaling initiated by neurons.

To further test the idea that rearing-dependent plasticity of TTCs does not depend on neuronal thermosensory cues, we quantified mutant *ttx-1*(p767) TPCs at 15°C, 20°C, and 25°C rearing conditions. The *ttx-1* gene encodes a transcription factor necessary for AFD neuron thermosensory fate, and its mutant genotype perceives temperature differently than normal (Fig. 4B). While TTCs for *ttx-1* mutants reared at 20°C and 25°C were shifted cooler compared to wildtype, the mutant nevertheless exhibited a similar pattern of rearing-dependent plasticity as wildtype N2 (rearing temperature differences for *ttx-1*(p767) at 32.5°C F_2,139_=9.98, P<0.0001; Fig. 4B). This observation reinforces our conclusions from the *trpa-1* mutant regarding the notion that neural signaling, and AFD thermosensation in particular, is not involved in controlling plasticity of behavioral TTCs in response to rearing conditions despite the fact that neural control contributes to non-rearing-dependent TTC shape.

## Discussion

We designed and implemented a micro-droplet swimming assay of nematode locomotory behavior to characterize the environmental plasticity and genetic modulation of behavioral reaction norms. These multidimensional data yield phenotypic functional responses that take the form of thermal tolerance curves (TTC) to capture distinct thermal performance profiles among natural isolates, mutant alleles, and rearing conditions. By screening TTCs for 58 *C. elegans* mutant strains with known or suspected roles in sensory perception and locomotion, we found potential neural controls that contribute to TTC shape. Mutations to thermosensory pathways are capable of shifting the decision-making process to cause animals to continue swimming at higher temperatures or to cease moving at lower temperatures than wild-type worms. These experiments provide evidence for a neural “decision” to stop locomotion toward the upper portions of thermal tolerance curves, in addition to physiological limits of high temperature stress on locomotory activity. In addition to neural components, we also show that *trp-1* expression in non-neural tissue affects the plasticity of the TTC.

Previous work on thermotaxis (15, 45) and thermal avoidance (46) has shown that *C. elegans* thermal behavior can be modified by experience. In a similar fashion, through behavior, the animal can “choose” to not perform at its peak, and the less-than-peak performance might be optimal for the organism. In this way, behavioral TTCs in our experiments differ in part from “classic” thermal performance curves of maximum sprint speed or fecundity that test absolute physiological thermal limits of the organism (12, 21). Even with neural control over temperature-dependent behavior, however, disrupted homeostasis must also contribute to TTC profiles at extreme temperatures. Nevertheless, our observations from sensory gene mutants that swim actively at temperatures even more extreme than wildtype argue that hard cell physiological limitation is not the sole determinant of locomotory declines in *C. elegans*.

Reduced locomotory activity of *C. elegans* from elevated temperature depends on continuous exposure to temperatures above a threshold near 30°C and exposure to just 80 s of extreme temperatures >39°C can alter locomotory behavior for at least an hour. Moreover, plasticity in animal TTC profiles derives from differences in age and, most importantly, the temperature of rearing conditions. We demonstrated that this TTC plasticity due to rearing conditions also has a genetic basis, such that knockout of the *trpa-1* ion channel ablates the normal sensitivity that adult *C. elegans* have to the rearing temperature conditions they experienced as larvae, resulting in a ‘canalized’ behavioral response.

Strikingly, it is a non-neural component of thermal tolerance regulation that mediates the *trpa-1*-dependent sensitivity of TTC profiles to rearing conditions. Previous work demonstrated that cold sensing of the *C. elegans* intestine regulates lifespan (42), and here we show that intestinal expression of *trpa-1* affects sensorimotor responses such as thermal tolerance. Thus, normal thermal response behaviors depend crucially on integration and feedback between distinct tissue types, including neurons that contribute to TTC shape and intestinal cells that contribute to how the animal assimilates ‘memory’ of rearing conditions. Other non-neural tissues also are reported to affect thermal behavior in *C. elegans*, with even sperm cells shown to influence thermal performance of the organism (47). In addition, hormone signals from many other tissues affect general worm locomotory behavior, including insulin or steroidal signaling (48, 49). It is clear that we are only beginning to understand how the whole body of *C. elegans* is integrated to regulate its sensory behavior.

Viewed through the lens of adaptive phenotypic plasticity (50), the “decision” to stop moving at high temperatures likely reflects some kind of preventative or protective response to improve fitness. Indeed, studies showing that mutants with less quiescence after exposure to high temperature have lower survival rates suggests that cessation of locomotion may be important for recovery from or coping with thermal stress (22). Given that normal functioning of *trpa-1* confers TTC plasticity to rearing conditions, the extent to which the sensitivity of TTCs to rearing conditions are adaptive would imply that *trpa-1* represents an “adaptive plasticity” gene. Future work that connects behavior to fitness of the *trpa-1* mutants at different temperatures, including low temperatures, would also help determine whether the plasticity that gets ablated by knockout of *trpa-1* is related to any adaptive role for *trpa-1* signaling.

## Conclusion

Understanding the environmental plasticity and genetic evolvability of the thermal ecology for ectothermic organisms is fundamental, given how temperature exerts profound effects on all levels of biological function with special urgency due to the effects of recent climate change. Here we developed an assay to quantify the thermal tolerance of *C. elegans* across an ecologically relevant temperature range. Through a canvasing of candidate genes, we show that the thermal performance of *C. elegans* is sensitive to genetic perturbations of both behavioral decision-making and physiological limits.

## Materials and Methods

### Micro-droplet assay of thermal behavioral responses

We constructed an experimental apparatus to quantify nematode locomotory behavior (Fig. S1) across a wide range of temperature. The device consists of a high-resolution video camera (Allied vision technology, model GX3300 with Nikon AF-Nikkor 80mm lens) mounted above a temperature-controlled aluminum stage (165mm x 58mm x 4.5mm). The imaging stage is obliquely illuminated by two LED strips. Two thermoelectric control devices (TEC), cooled by liquid CPU coolers (Swiftech, MCW30; Thermo Scientific, NESLAB Digital One RTE7), control the temperature of the stage. A custom-built closed-loop input/output controller, driven by a program written in LabVIEW (National Instruments, Texas) controls the TECs, and a thermocouple (±0.1°C) measures the stage temperature near the stage center.

To assess nematode behavior on the instrument stage, we placed individual worms within ∼2 µL NGM liquid droplets on a Teflon (PTFE) printed microscope slide that hydrophobically constrains the droplets in a 2D array (Tekdon slide ID: 24-20). Worms were distributed in the middle 3 x 6 wells of a slide patterned with a 3 x 8 array of 4mm diameter wells. The slide was sealed with a coverslip that contacted the NGM droplets, using two layers of double-sided sticky tape and M10 Apiezon vacuum grease along its edges.

### Experimental protocol for behavioral quantification

Using a worm pick, worms from stock plates were placed into NGM buffer at room temperature, swirled to wash off bacteria and poorly swimming worms were removed. We pipetted 1.8 µL of NGM buffer containing a single worm into each of 18 wells per slide (∼ 7 min total prep time), with each half of the slide holding a randomized pair of strains for experiments comparing different genotypes.

Our standard experimental assay comprised of sixteen, ∼1°C temperature steps, of 80 sec duration, where we let the system to come to equilibrium for 60 seconds and then captured 20 seconds of behavioral video at 15fps (3296 x 1600 pixels). Image capture and primary analysis used custom software written with LabVIEW. For each video frame, the analysis program identifies the worm in each droplet using particle detection and measures its area and center-of-mass position. Worm size and swimming speed were used as criteria for removing data affected by poor worm tracking or abnormally moving worms. Cutoffs include a maximum allowable worm size (mean area 375 pixels, maximum area 803 pixels), a maximum percentage change in worm size (31% between adjacent frames threshold value) and a maximum change in worm position between frames (15 ‘pixel’ width, threshold value), determined from a pilot study of 198 animals from two *C. briggsae* strains (AF16 and HK104; similar size and behavior as *C. elegans*). Each temperature step recording required at least 100 of the possible 300 frames of data in order to be included in downstream analysis.

We define a “Locomotion Index” (LI) to quantify movement activity by comparing worm images between video frames. The LI is calculated as the ratio of the number of non-overlapping worm pixels between two frames over the total number of worm pixels for the two frames. When a worm moves slowly between frames, nearly all pixels will overlap and the LI will be close to 0. Fast worm movement results in less pixel overlap and the LI will approach a maximum value of 1. In practice, stochastic pixel noise produces a non-zero lower bound of LI values (∼0.05) and the partial overlap of worm position between frames yields an upper bound (∼0.60). We did not normalize LI within strains in order to capture differences in the baseline. We indicate the sampling rate of the LI as LI#, where # indicates the index of the second frame used to calculate the LI (e.g. LI1 indicates successive frames, while LI7 indicates every 7^th^frame). Both LI1 and LI7 measurements gave qualitatively similar results, but we chose one over the other depending on how slow the animals were moving. We determined whether animals were moving or not based on a threshold of the mean locomotion index calculated seven frames apart (LI7, 0.47 second resolution). Only 2.5% of worms exceeded the threshold value at extreme temperatures (LI7 > 0.14), when all worms are paralyzed, using the same dataset used to calculate quality control cut-offs.

We determined statistical significance of differences between strain performance with one-way ANOVA and post-hoc tests at particular temperatures in the thermal response. ANOVA post-hoc tests used either Dunnett’s test with N2 as control (51) or Tukey-Kramer Honestly Significant Differences (HSD). In comparisons of plasticity for *trpa-1* experiments, we also tested for overlap of 95% confidence intervals for parameters from a three parameter logistic function fit to LI using non-linear model fitting in JMP 10.0.0: 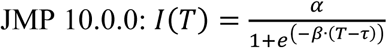, where *T* is droplet temperature and τ is the inflection point temperature in the TTC.

### Worm strains and preparation

We reared 58 isogenic strains each carrying a homozygous mutationplus two “wildtype” strains of *C. elegans* using standard protocols (52) (Supplementary Table T1), obtained from the *Caenorhabditis* Genetics Centre (University of Minnesota, Minneapolis, MN, USA) which is funded by NIH Office of Research Infrastructure Programs (P40 OD010440) with some mutants created by the *C. elegans* knockout consortium (53). We selected strains based on published literature that indicated roles in thermotaxis, thermosensation, mechanosensation, chemosensation, locomotion, and neural function. All mutant strains derive from the N2 genetic background, so N2 represents the primary control for comparison. We assayed two common wild-derived *C. elegans* strains as controls: the archetypal N2 strain (Bristol, England) and the genetically distinct CB4856 (Hawaii, USA), which have been used in QTL mapping of other temperature-dependent traits and behaviors (54-58). These strains provide a useful reference for the potential natural range in phenotype without disruptive gene mutations.Behavior assays used only well-fed adult worms, except in experiments with dauer, L3 and L4 stage larvae. For rearing temperature manipulations, we raised all worms from eggs at the desired temperature from parents raised at the same temperature.

## Acknowledgements

We thank Jiwon Shih for preliminary work on the droplet assay. Some strains were provided by the CGC, which is funded by NIH Office of Research Infrastructure Programs (P40 0D010440).

## Availability of data and materials

All data analyzed for this paper are available from the corresponding authors on reasonable request.

## Authors’ contributions

GWS, ADC, and WSR wrote the manuscript and designed the experiments. GWS and DM performed the experiments. GWS performed the data analysis. GWS and ADC performed the statistical testing. All authors read and approved of the manuscript.

## Ethics approval and consent to participate

Not applicable.

## Consent for publication

Not applicable.

## Supplementary figures

**Supplementary Figure 1.**
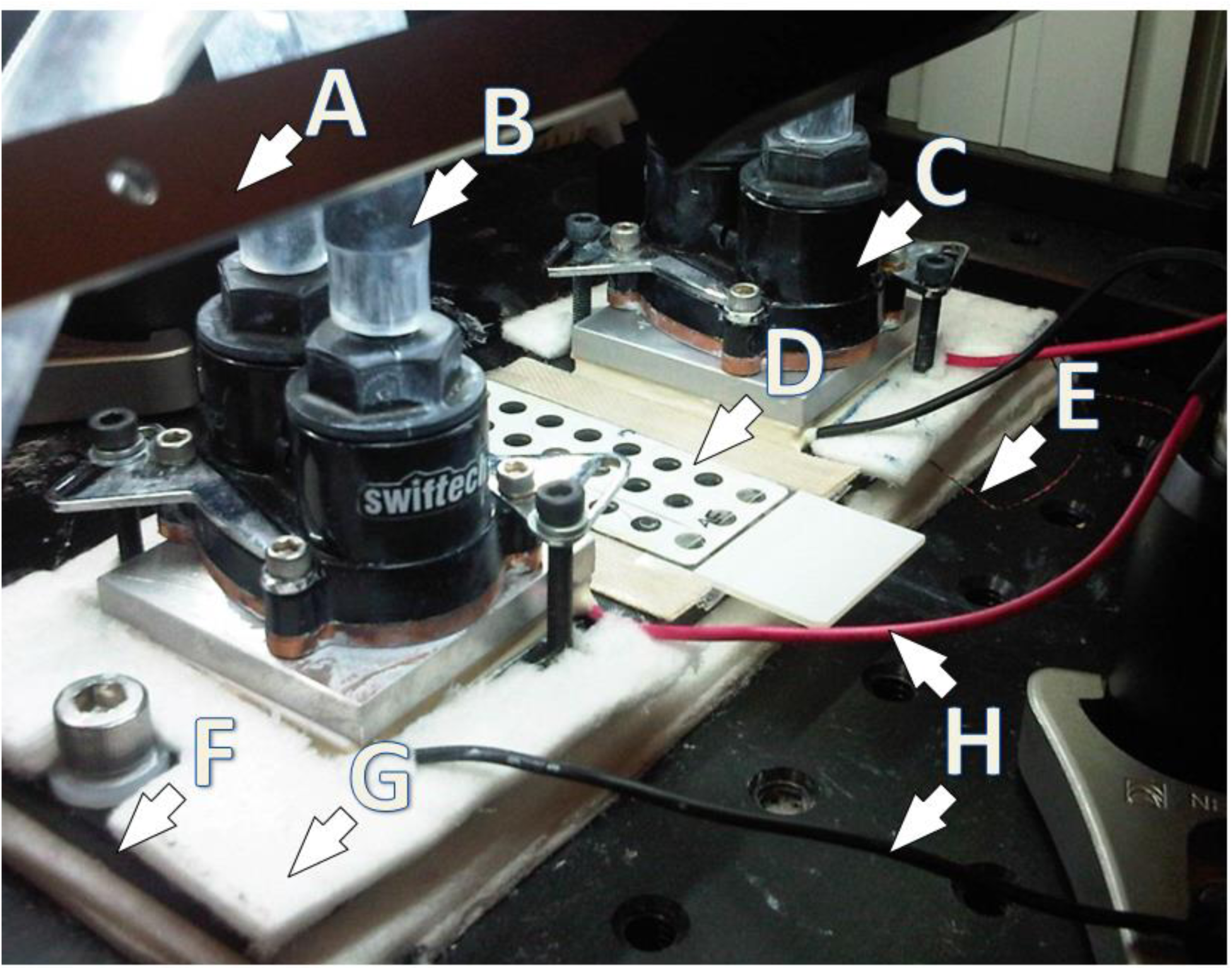
Droplet apparatus and setup. (A) LED light strip (B) water tube to temperature-controlled water bath (C) water block connected above thermo-electric cooler (TEC) pad (D) droplet slide with droplets and cover slip (E) thermocouple wires for detecting temperature of aluminum block (F) edge of aluminum block/stage (G) fiberglass insulation around aluminum block (H) wires from TECs to I/O controller. Not pictured are the I/O controller and the camera mounted 24 cm above the stage. For scale, each circular well at (D) is 4mm in diameter and TECs are 45mm apart.

**Supplementary Figure S2.**
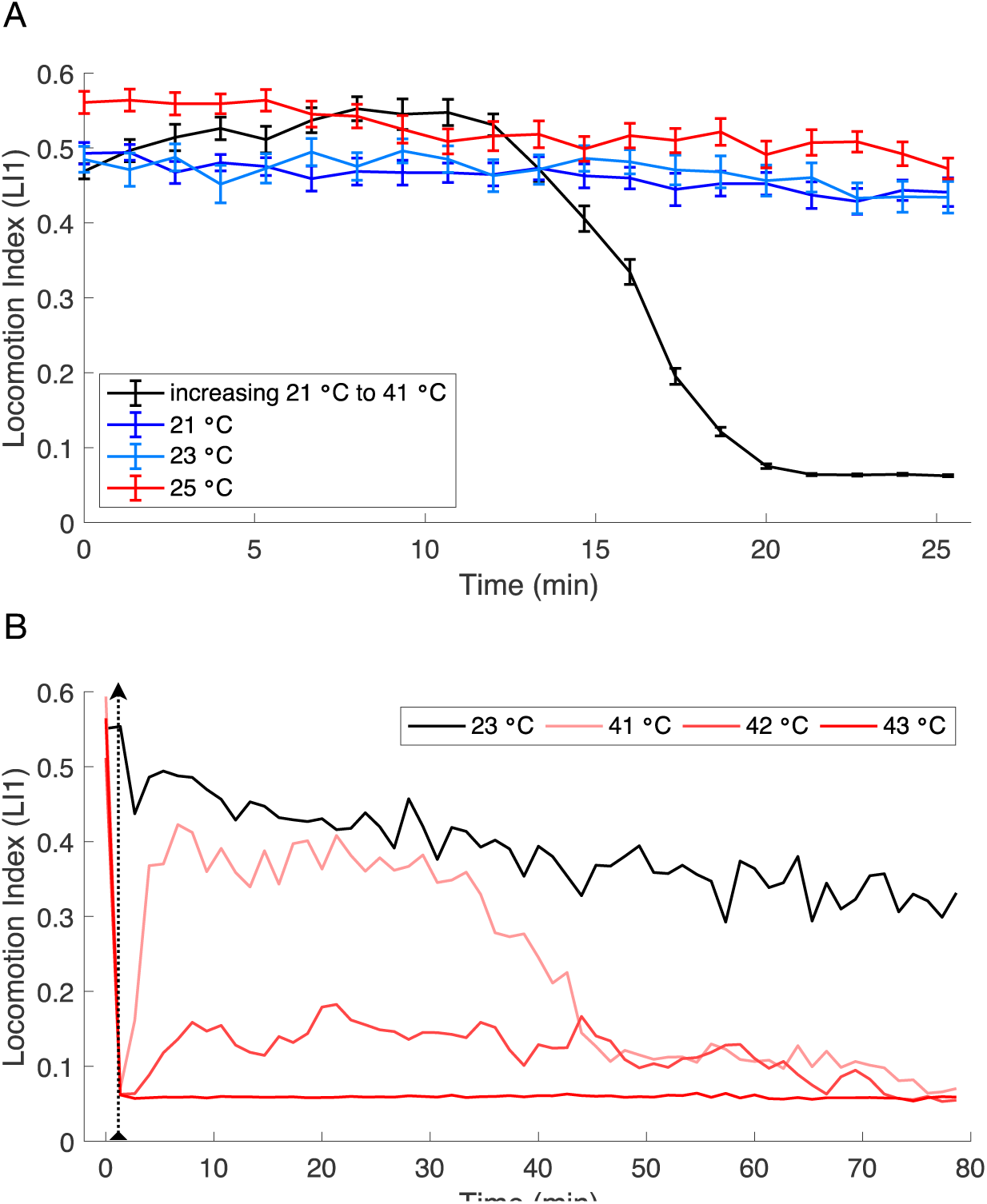
Locomotion Index. (A) Locomotion Index at constant temperatures. LI1 is nearly constant for the duration of the experiment (21 min.) at benign temperatures. (B) Longer term (58 steps ∼ 77min) effects of brief high-temperature exposure. High temperature exposure of 80 sec occurs at dotted line. Control worms held at a constant 23°C (black line) for the whole experiment gradually slowed their locomotion to approximately 60% of the level seen at the start of the experiment. This slowing of average locomotion occurs because a fraction of worms stop or entering cycles of episodic swimming rather than from all individuals slowing down. *n*=18 worms tested for each condition, minimum of 8 worms included in means. Error bars are ± SEM.

**Suplementary Figure S3.**
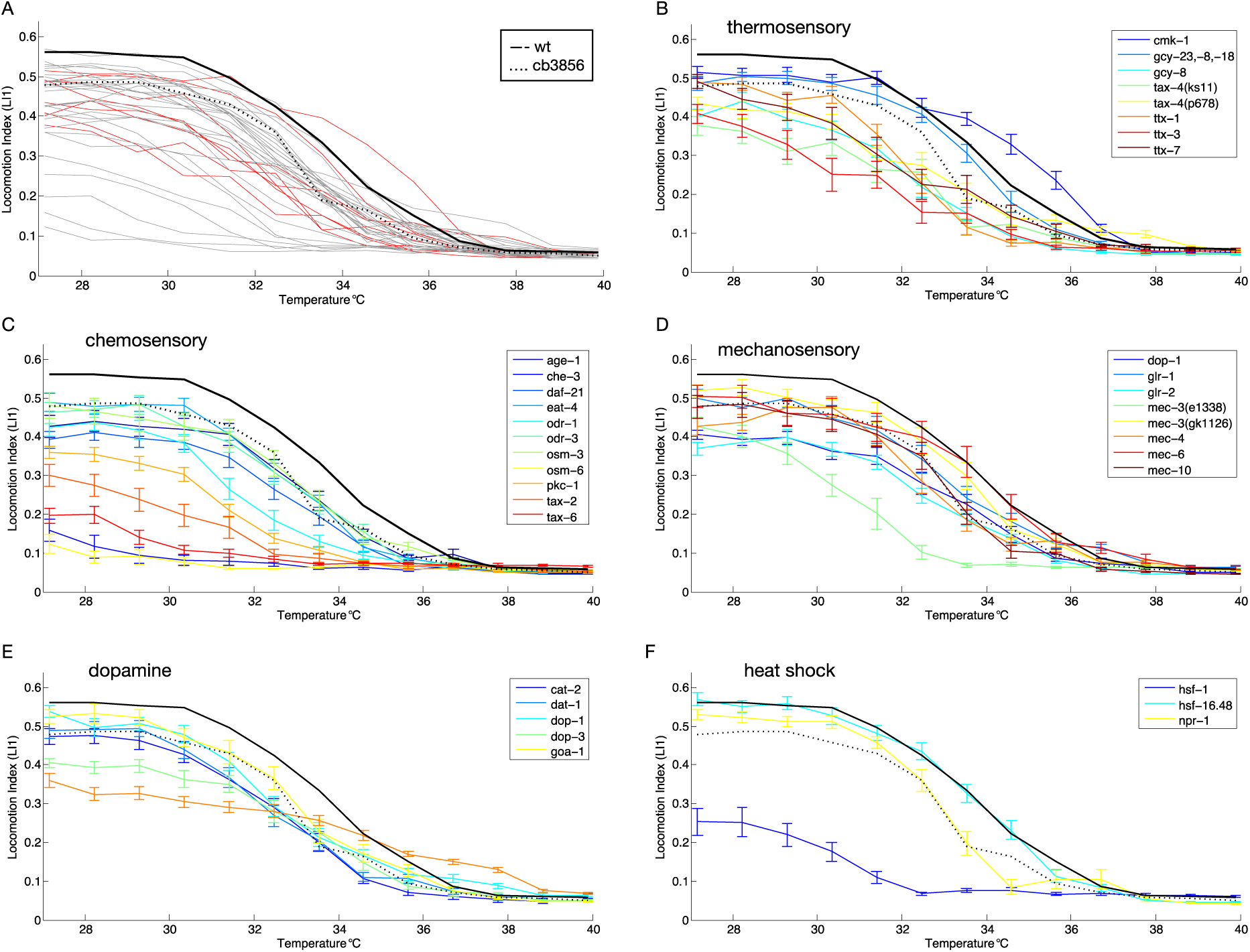
(A) Thermal reaction norms of swimming behavior for 58 *C. elegans* genetic mutant strains (gray and red lines) and two wildtype strains (black line = N2, dashed line = CB4856). All worms reared at 23°C; 17 −36 individuals included in calculations at each assay step for each strain (Supplementary Table S1). TPCs for subsets of mutant strains with sensory disruptions from (A) are shown for thermosensory defects (B), chemosensory defects (C), and mechanosensory defects (D). (E) TPCs for strains with gene mutants affecting dopamine signaling. (F) TPCs for strains with mutations in genes involved in heat shock response, oxygen sensing, and muscle activity. Error bars in (B-F) indicate ± SEM; TPCs for wildtype strains N2 and CB4856 are shown in all of (A-F) for reference.

**Supplementary Figure S4.**
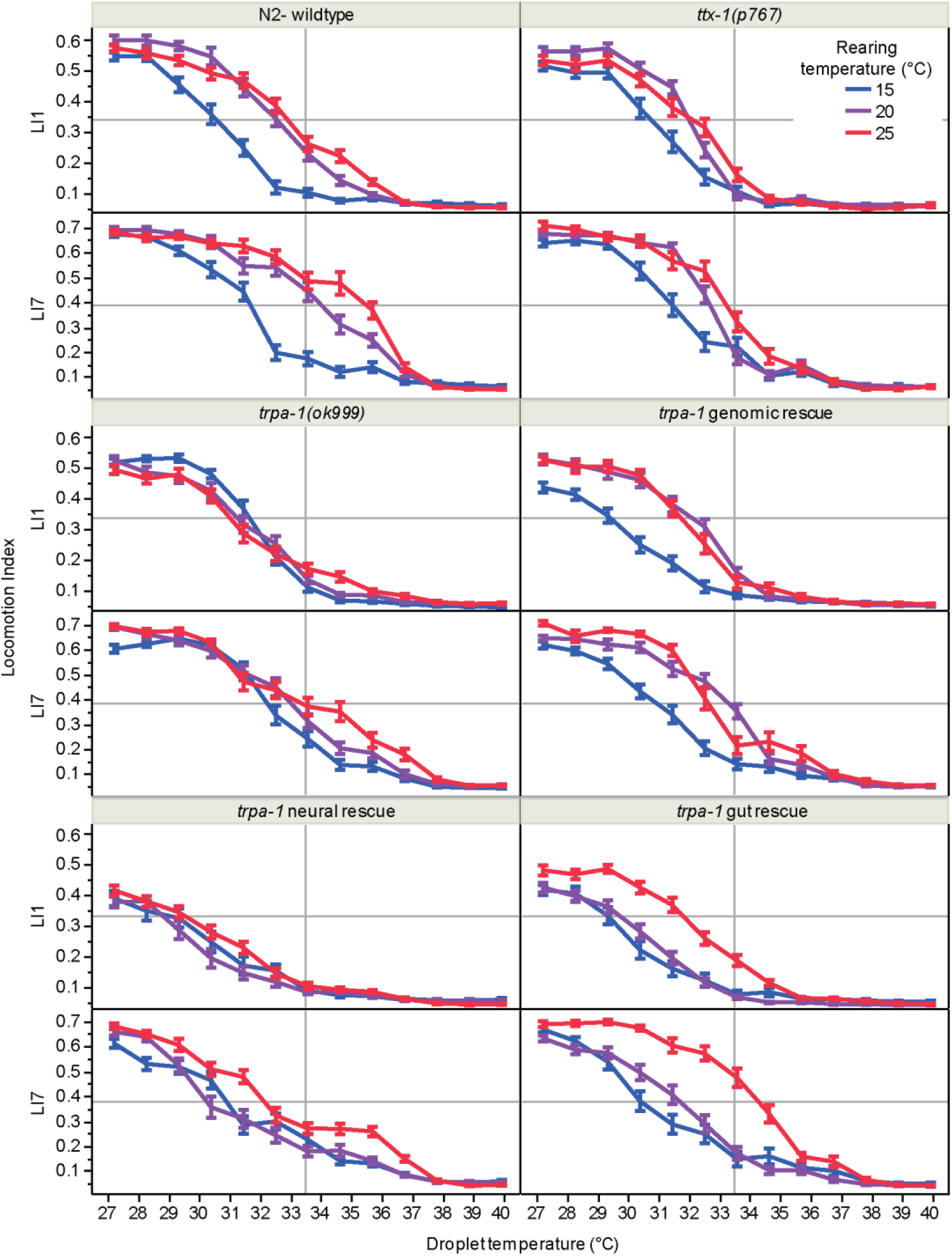
Plasticity of thermal performance response to rearing temperature of tissue specific *trpa-1* rescues.

**Supplementary Figure S5.**
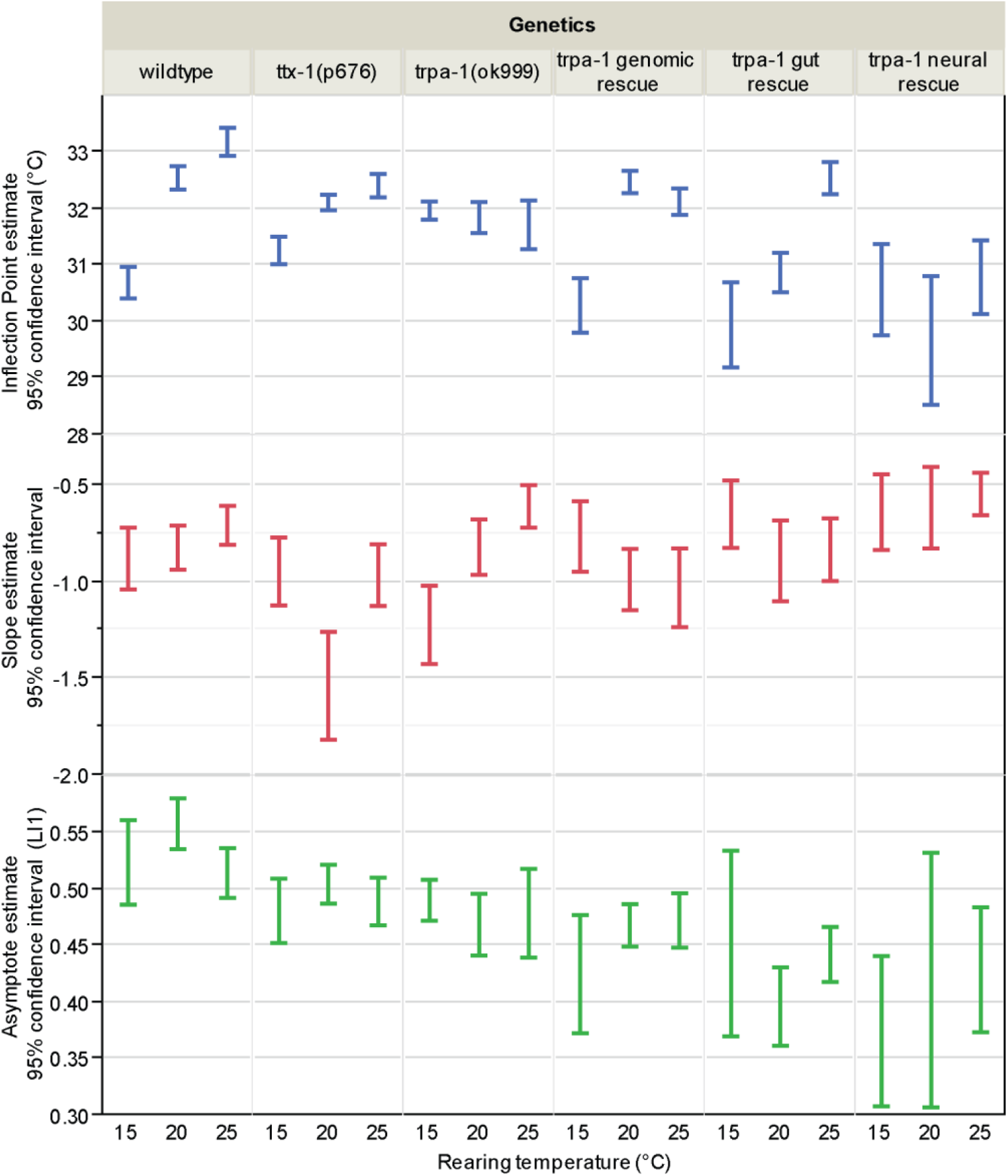
Function fit parameter estimate 95% confidence intervals for *trpa-1* rescue experiment. 95% confidence intervals for three parameter logarithmic function fit to LI1 data for strains in *trpa-1* rescue experiments at three rearing temperatures.

**Supplementary Table S1.** Mutant strains quantified for differences in temperature-dependent locomotory behavior.

| Strain | Mutant gene(s) | Allele | Mutation type* |
| --- | --- | --- | --- |
| N2 | wildtype Bristol |  |  |
| CB4856 | wildtype Hawaii |  |  |
| TJ1052 | age-1 | hx546 | substitution |
| CB6055 | bus-8 | e2698 | substitution |
| CB1112 | cat-2 | e1112 | substitution |
| TN101 | cha-1 | cn101 | from EMS |
| CB1124 | che-3 | e1124 | substitution |
| PY1589 | cmk-1 | oy21 | substitution |
| PR673 | daf-21 | p673 | from EMS |
| RM2702 | dat-1 | ok157 | from UV/TMP |
| JT748 | dgk-1 | sa748 | from EMS |
| LX636 | dop-1 | vs101 | deletion |
| LX703 | dop-3 | vs106 | deletion in promotor and 1st transmembrane domain |
| BZ33 | dys-1 | eg33 | from EMS |
| DA572 | eat-4 | ad572 | from X-ray |
| MT1212 | egl-19 | n582 | substitution |
| VC461 | egl-3 | gk238 | deletion |
| IK597 | gcy-23; gcy-8; gcy-18 | nj37, oy44, nj38 | all deletions |
| IK800 | gcy-8 | oy44 | deletion |
| KP4 | glr-1 | n2461 | substitution |
| RB1808 | glr-2 | ok2342 | deletion |
| JT734 | goa-1 | sa734 | substitution |
| PS3551 | hsf-1 | sy441 | substitution |
| RB791 | hsp-16.48 | ok577 | deletion |
| MT382 | lin-11 | n382 | from EMS |
| CB1515 | mec-10 | e1515 | substitution |
| CB1338 | mec-3(e1338) | e1338 | Insertion - frameshift |
| VC2396 | mec-3(gk1126) | gk1126 | insertion/deletion |
| CB1339 | mec-4 | e1339 | substitution |
| CB1472 | mec-6 | e1342 | substitution |
| DA609 | npr-1 | ad609 | substitution |
| CX2065 | odr-1 | n1936 | from EMS |
| CX2205 | odr-3 | n2150 | from EMS |
| MT3631 | osm-3 | n1545 or n1540 | from EMS |
| PR811 | osm-6 | p811 | substitution |
| IK105 | pkc-1 | nj1 | substitution |
| PR671 | tax-2 | p671 | substitution |
| FK129 | tax-4(ks11) | ks11 | substitution |
| PR678 | tax-4(p678) | p678 | substitution |
| PR675 | tax-6 | p675 | substitution |
| GR1321 | tph-1; cam-1 | mg280 & vs166 | deletion |
| VC160 | trp-1 | ok323 | deletion |
| RB1052 | trpa-1 | ok999 | uncurated |
| PR767 | ttx-1 | p767 | substitution |
| FK134 | ttx-3 | ks5 | substitution |
| IK575 | ttx-7 | nj40 | substitution |
| TN110 | twk-18 | cn110 | substitution |
| MT1684 | unc-105(n490n785) | n490n785 | double mutation, |
| MT1098 | unc-105(n506) | n506 | uncurated |
| VC854 | unc-2 | gk366 | deletion |
| BC18 | unc-22 | s13 | from EMS |
| CB403 | unc-29 | e403 | substitution |
| CB251 | unc-36 | e251 | substitution |
| ZZ20 | unc-38 | x20 | uncurated |
| CB177 | unc-46 | e177 | substitution |
| CB382 | unc-49 | e382 | substitution |
| BC347 | unc-54 | s74 | substitution |
| CB1068 | unc-79 | e1068 | from EMS |
| MT1085 | unc-8 | n491 | substitution |
| CB1069 | unc-80 | e1069 | from EMS |
| MT200 | unc-93 | n200 | from EMS |

\*Mutation information from wormbase.org

